# sv-channels: filtering genomic deletions using one-dimensional convolutional neural networks

**DOI:** 10.1101/2024.10.17.618894

**Authors:** Luca Santuari, Sonja Georgievska, Arnold Kuzniar, Brent S. Pedersen, Carl Shneider, Sarah Mehrem, Lars Ridder, Wigard P. Kloosterman, Jeroen de Ridder

## Abstract

Structural variant (SV) detection in human genomes using short-read sequencing data is hindered by false positives, arising from sequencing and mapping artifacts that mimic genuine SV signals. Despite advances, state-of-the-art SV callers like GRIDSS and Manta exhibit trade-offs between precision and recall, with GRIDSS offering the highest precision and Manta excelling in recall. To address these limitations, we introduce sv-channels, a novel deep learning model designed to improve the precision of SV detection by leveraging read information at call sites. Our method effectively reduces false positives in Manta’s deletion callsets, achieving precision that surpasses GRIDSS while maintaining a recall rate comparable to Manta. This represents a significant improvement in SV detection, leveraging Manta’s high recall through deep learning and paving the way for more accurate genomic analyses. The sv-channels codebase is openly accessible on GitHub at https://github.com/GooglingTheCancerGenome/sv-channels enabling further research and application in the field.

## Introduction

Structural variants (SVs) are genomic rearrangements defined as 50 base pairs (bp) or longer that can either be balanced (inversions, intra-, and inter-chromosomal translocations) or unbalanced (deletions, novel sequence insertions, tandem, and interspersed duplications) (Ho et al., 2020). SVs account for the largest sequence diversity across individuals (Auton et al., 2015) and have been linked to genetic diseases like neurodevelopmental disorders (D’haene and Vergult, 2021) and cancer (Li et al., 2020). SVs can have profound effects on the genomic landscape. They can lead to gene copy number changes or gene fusions, and they can rewire the gene regulatory code by modifying the local three-dimensional conformation of a genomic region, disrupting enhancer-gene interactions, or establishing new ones (Li et al., 2020). Characterizing how SVs impact DNA is important on multiple levels: to understand the etiology of a disease, track disease progression, and design novel targeted therapies. A pre-requisite for this is the comprehensive and accurate identification of the set of SVs in a genome. To date, Illumina paired-end sequencing remains the technology of choice for variant detection in large-scale studies due to its cost-effectiveness, high base throughput and quality, and widespread adoption. SV detection has relied on heuristic algorithms, SV callers, that look for multiple signatures in alignment data where reads originating from a donor sample have been aligned to a reference sequence to find SV breakpoints. These breakpoints are defined as positions that are adjacent in the donor DNA but not in the reference. SV signatures include variable read depth of coverage (RD), clipped reads (CR, when reads only partially match the reference sequence and are truncated), split reads (SR, when the two parts of a read match two different locations in the reference), discordant pairs (DP, read pairs with abnormal distance and orientation) and open-end anchored reads (OEA, pairs where only one of the read is mapped). Among SV callers, two in particular have been shown to have consistently high performance on Illumina data (Cameron et al., 2019): GRIDSS (Cameron et al., 2017, 2021) and Manta (Chen et al., 2016). GRIDSS collects CR, SR, DP, OEA and reads with indels, and uses a positional de Bruijn assembler to construct contigs that are used to resolve SV breakpoints. Manta uses SR and DP evidence to build a breakend graph to identify the SVs and performs targeted assembly only as a post-processing step to refine the variants.

We noted that Manta has high recall, but lower precision than GRIDSS (Cameron et al., 2019). Therefore, we set out to develop an approach that could remove false positives from Manta’s SV callsets thereby increasing its precision to levels comparable to GRIDSS while maintaining its high recall. Deep Learning (DL) models have revolutionized many fields of biology including genomics and proteomics. DL models have been used to predict DNA accessibility from sequence (Kelley et al., 2016), improve gene expression prediction from DNA sequences (Avsec et al., 2021), and protein sequence-to-structure prediction (Jumper et al., 2021). Small variant calling has seen DeepVariant (Poplin et al., 2018) outperform competitors using the Inception_v3 architecture (Szegedy et al., 2015) (24 millions trainable parameters) to classify images of read alignments at variant sites into true variants or sites that match the reference sequence. DeepSV (Cai et al., 2019) builds on the idea of DeepVariant to classify genomic regions with deletions longer than 50 bp after encoding them into image representations (2D arrays). Samplot-ML (Belyeu et al., 2021) and DeepSVFilter (Liu et al., 2021) are two additional examples of SV filtering methods for short read data where a CNN-based classifier is applied to visual representations of candidate genomic regions for SVs. It remains open whether a simpler representation could be sufficient for a deep learning model to recognize true SVs. For instance, encoding SV signals as one-dimensional (1D) arrays instead of images (2D arrays) would simplify the type of model needed, which in turn would require less time for model optimization and training. It would also lead to a reduction in the size of the training data. Germline SVs only account for a few thousand variants per genome, far from the order of millions reached by short variants (SNVs and InDels). This puts a constraint on the complexity of the model architecture that needs to be much simpler than the model used in DeepVariant to avoid overfitting. To test whether a one-dimensional representation of SV signals could be used to improve the precision of Manta’s callsets, we created sv-channels, a novel deep learning approach that uses one-dimensional convolutional neural networks to classify SVs called by Manta as true SVs or false positive calls. We tested it on deletions that represents the most abundant class of SVs. Our model can effectively remove false positive DEL calls from Manta callsets, resulting in a large gain in precision (82.87% when tested on the 1KG sample HG00420), while keeping a high recall.

## Methods

### sv-channels methodology

We propose a deep learning model that is trained to retain true positive calls from SV callsets obtained with the SV caller Manta. SV signals are encoded in one-dimensional arrays called ‘channels’ (Figure 1 A). The first dimension in a channel represents genomic position relative to a putative breakpoint. Channels are stacked into 2D arrays and sliced at intervals [breakpoint1 - window_size/2, breakpoint1 + window_size/2] and [breakpoint2 - window_size/2, breakpoint2 + window_size/2], where breakpoint1 and breakpoint2 are the first and the second breakpoint of the SV and window_size is a parameter that depends on the fragment length of the sample. This results in two 2D SV-arrays. SV-arrays are joined into a 2D array of dimension [2^*^window_size+buffer_size, 35] where window_size is the length of each interval, buffer_size is the size of the 2D arrays of zeros inserted between the two windows to avoid artifacts when the convolutional filter passes along the arrays, and 35 is the number of channels. We call this 2D array “window-pair”. At this point, channels dependent on either read pairs or split reads that bridge both intervals are added to the windows-pair (see Supplemental Table 2 for the complete list of channels) along with other signals within and between reads. The channels encode qualitative and quantitative information of both the reference sequence and the reads aligned at the two intervals centered on the SV breakpoint positions. For split reads and discordant reads there are channels encoding the read orientation, whether a read is split on its right or on its left and whether both parts of a read pair or a split read are falling into the window-pair intervals or only one (orphan parts). For split reads we record the information at the split position, while for discordant reads we use the right-most position of the left read and the left-most position of the right read. We also consider whether a read has an abnormally high number of mismatches, if it has a small deletion or insertion encoded in the CIGAR string, and if it has a mapping quality lower than a certain threshold. Window-pairs are labeled based on the overlap of the two intervals with DEL calls from a high-quality truth set, either with the “DEL” for deletions supported by the truth set or “noDEL” for false positive calls (Figure 1 B). Labeled windows-pairs form the training set are used to train a convolutional neural network with multiple convolutional layers, one dense layer, and a final Soft-Max layer. Hyperparameters (regularization rate, initial learning rate, number of nodes in the convolutional layers and in the dense layer, number and size of the convolutional filters, and the rate rate of dropout) are optimized using sequential Bayesian optimization with Gaussian Processes (using the gp_optimize function included in the Python library scikit-optimize (Head et al., 2018)). sv-channels is implemented in Python as a command-line tool (Santuari et al., 2023). Notably, the code has been optimized to allow easy and fast creation of channels from the read alignment files in BAM format, thus maximizing the utility of the tool. The analysis of an input BAM file of a sample results in a VCF file with SV calls (DELs only) that include the posterior probabilities returned by the sv-channels model as SV qualities (QUAL). Instructions on how to run sv-channels are included in the GitHub repository: https://github.com/GooglingTheCancerGenome/sv-channels.

**Figure 1.**
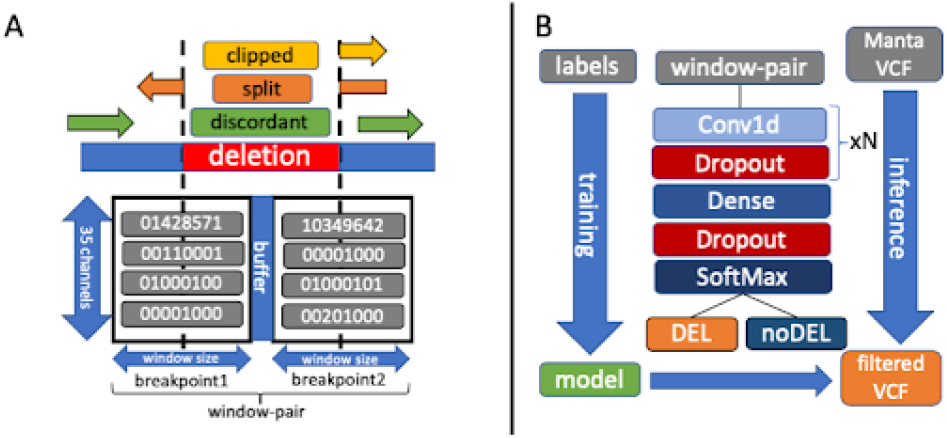
sv-channels input data and architecture. (**A**) Channels are extracted from genomic intervals centered on candidate deletion breakpoints. SV signals are encoded into one-dimensional arrays called channels. Channels are stacked into 2D arrays called window-pairs, with zero padding in between. As an example, the channel for soft-clipped reads will increment the relative base position each time a read with a soft-clipped read is seen. (**B**) Labelled window-pairs are used to train a convolutional neural network to classify Manta deletions calls into either true deletions or false positives. Model hyperparameters are optimized in the inner loop of a nested cross-validation procedure using sequential Bayesian optimization with Gaussian Processes.

### Simulation of structural variants

To test sv-channels on simulated SVs we implemented sv-gen (Kuzniar and Santuari, 2023), a snakemake (Köster and Rahmann, 2012) workflow to generate artificial read alignment data with SVs introduced at random positions. We simulated 2,000 SVs of each type (deletions, insertions, inversions, duplications and inter-chromosomal translocations), with size ranging from 50 bp to 5,000 bp into chromosome 10 and chromosome 12 of the 1000 Genomes human reference genome (hs37d5). We simulated both homozygous and heterozygous SVs. From the chromosomes containing the SVs we generated read alignment data with variable coverage (5x, 10x, 15x, 30x, 45x, 60x, 75x, 90x; with fixed insert-size at 500 bp and read-length at 150 bp), insert-size (200 bp, 250 bp, 300 bp, 400 bp, 500 bp, 600 bp; with fixed coverage at 30x and read-length at 150 bp) and read-length (36 bp, 50 bp, 75 bp, 100 bp, 150 bp, 250 bp; with fixed coverage at 30x and insert-size at 500 bp). SVs were called with Manta (version 1.1.0) and Manta DELs were filtered with sv-channels by retaining only calls with SV QUAL (sv-channels posterior probabilities) equal or greater than 0.5.

### Training on the 1KG dataset

To test sv-channels on real germline deletions we selected 8 samples from the high-coverage 1000 Genomes (1KG) dataset (Byrska-Bishop et al., 2022). We used 7 samples for training (HG01053, HG01114, HG01881, HG02018, HG02924, HG03992, NA06991) and we used sample HG00420 as a hold-out test set. 30% of the training set was used for validation and the hyperparameters of the model were optimized using sequential Bayesian optimization with Gaussian processes in the inner loop of a 10-fold nested cross-validation. We noted that data leakage between test and training set exists, as a result of germline deletions that are shared among samples (Figure 2). To test how the model would perform without this data leakage we used two cross-validation strategies. The first, called “Leave-One-Chromosome-and-Sample-out” (LOCaS) (Figure 3) is a nested cross-validation where in the outer loop iteratively one chromosome at a time is selected as test set, chromosome 22 is used as validation set and the remaining chromosomes form the training set. This is iterated such that all chromosomes apart from the validation chromosome are selected as a test set once. The inner loop is used to optimize the hyperparameters. The second strategy is a 2-fold LOCaS where in the outer cross-validation loop chromosome 22 is used as validation set and the remaining chromosomes are split into two sets; one is used as training set and the other is used as test set (Supplemental Figure 3 and Supplemental Figure 4). This is iterated across various splits of test and training set. In addition, we tested the LOCaS and 2-fold LOCaS strategies when, instead of using the whole chromosomes, each chromosome is divided into three roughly equal sets of DELs based on the position of the first breakpoint. In this case, instead of a full chromosome, one bin is considered at each time as test set in the Leave-One-Bin-and-Sample-out (LOBaS) cross-validation procedure (Supplemental Figure 5). Similarly, the 2-fold LOBaS procedure is similar to the 2-fold LOCaS procedure with bins that are used instead of whole chromosomes (Supplemental Figure 6).

**Figure 2.**
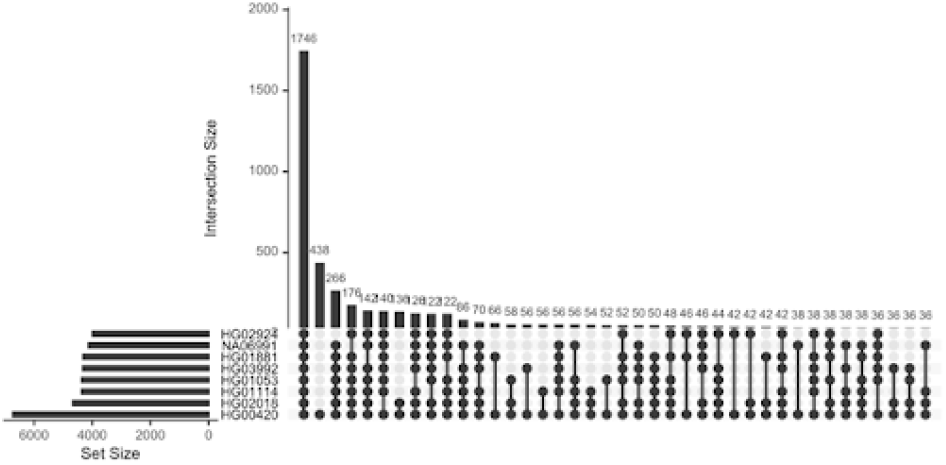
Overlap among deletions. Overlap among the truth set deletions of the 1KG samples considered in the study. The truth set for the sample HG00420 was used as a reference set to compute the overlap with the other truth sets.

**Figure 3.**
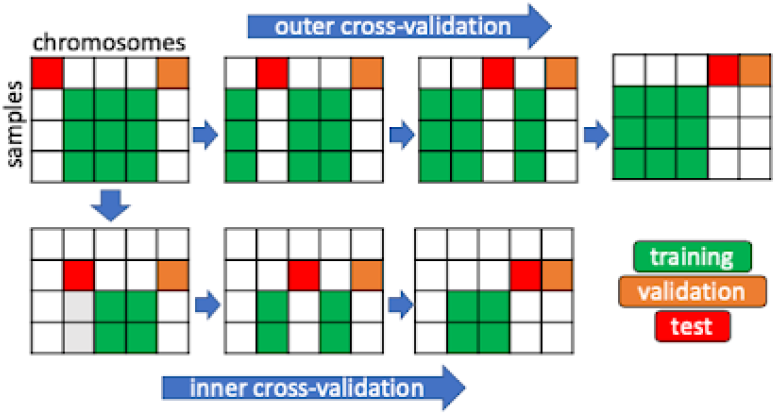
The cross-validation procedure. The nested Leave-One-Chromosome-and-Sample-out (LOCaS) cross-validation procedure with the inner and outer cross-validation highlighted. Chromosome 22 was used as validation set. The inner cross-validation is used to optimize the hyperparameters of the model.

## Results

### Performance on simulated data

We evaluated the performance of sv-channels varying read depth of coverage (coverage), insert size of the sequenced fragment (insert-size) and read length (read-length). Figure 4 summarizes the results, measured by the F1 score, defined as the harmonic mean of precision and recall. The largest decrease in F1 score can be seen at 5x coverage and for read-length equal to 75 bp. The lower performance of sv-channels at coverage less than 10 and read-length less than 100 bp suggests the importance of channels with SR information in making the prediction. Fewer or no split reads are expected to be present at DEL breakpoints when the coverage is as low as 5x or the read-length shorter than 100 bp.

**Figure 4.**
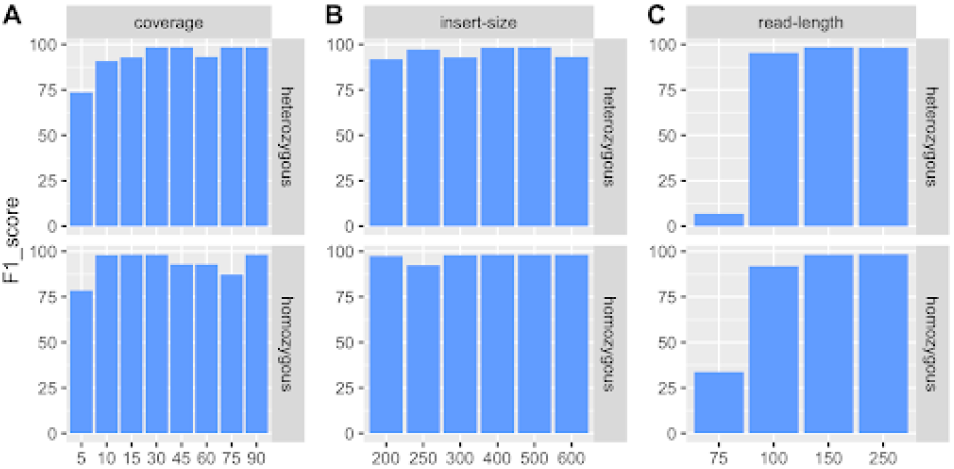
Evaluation on simulated data. F1 score for sv-channels when applied to simulated heterozygous (row 1) and homozygous (row 2) deletions. A) coverage B) insert size C) read length. Bars for reads of 36 bp and 50 bp are not shown because sv-channels did not produce a result. However, it should be noted that in current next-generation sequencing applications the minimum read length is usually 150 bp or longer.

### Performance on real data: 1KG dataset

sv-channels was applied to filter Manta’s deletions called on the sample HG00420 after training the model using seven samples (HG01053, HG01114, HG01881, HG02018, HG02924, HG03992, NA06991). This resulted in a 82.87% increase in precision and a 10.54% decrease in recall with respect to the unfiltered Manta callset (Figure 5). This represents a typical scenario where sv-channels would be trained on a set of samples and the trained model would be used to filter false positive deletions called with Manta on an unseen sample. However, this case leads to data leakage between the training and the test set, whereby the deletions commonly shared in the population are observed in both the training samples as well as the test sample. Most germline deletions that are common among the considered samples are shared by all of them (1746 deletions) (Figure 2). In practice this source of overtraining may be acceptable, as in a realistic application of sv-channels population SVs would also need to be detected. However, to evaluate if sv-channels can generalize beyond population SV signals and detect SVs de novo, we sought to assess its performance in the complete absence of data leakage using the LOCaS strategy applied to the HG00420 Manta DEL callset. This scenario led to a 74.08% increase in precision and a 15.91% decrease in recall with respect to the unfiltered Manta callset (Figure 5). The 2-fold LOCaS strategy (Supplemental Table 4) showed a 72.03% increase in precision and a 11.15% decrease in recall with respect to the unfiltered Manta DELs callset. When chromosome bins instead of whole chromosomes were considered, in the LOBaS strategy the model led to a 68.96% increase in precision and a 15.06% decrease in recall with respect to the unfiltered Manta callset (Supplemental Figure 5), while the 2-fold LOBaS approach led to a 64.98% increase in precision and a 10.16% decrease in recall with respect to the unfiltered Manta callset (Supplemental Figure 6). This shows that the model is still able to achieve good performance when only deletions that are unique to the training set are considered.

**Figure 5.**
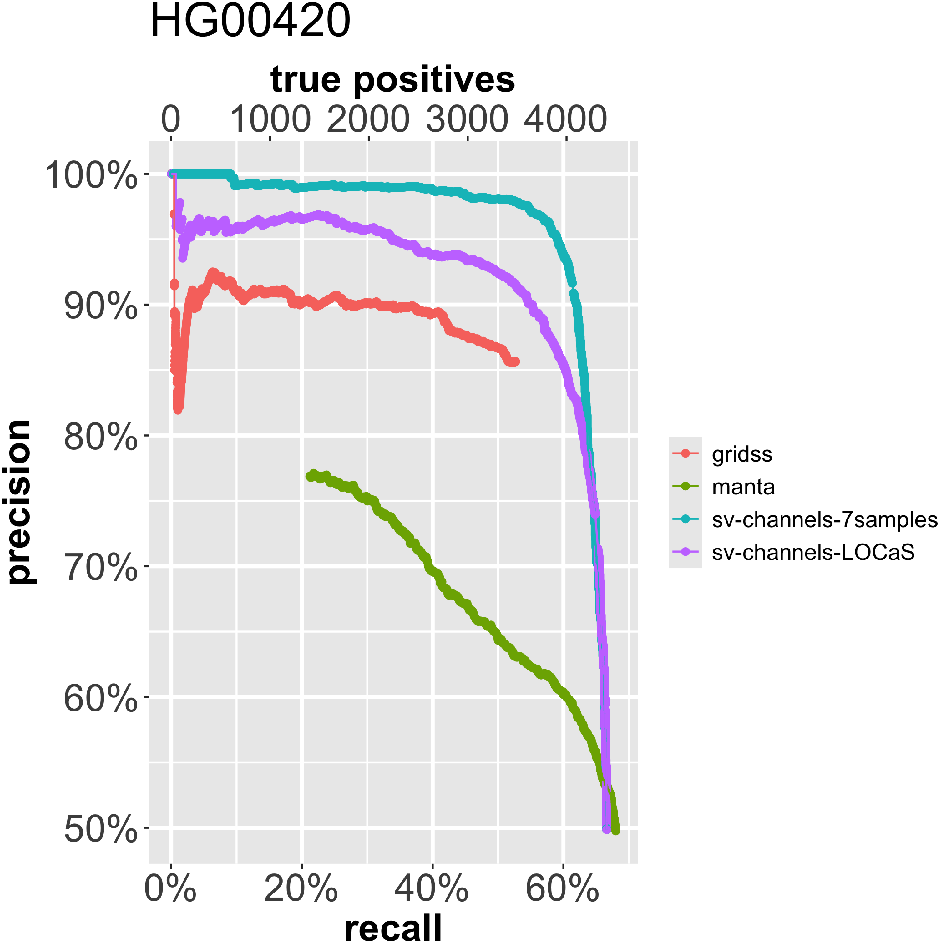
Evaluation on real data: HG00420. Precision-recall curve for GRIDSS, Manta and sv-channels on the 1KG sample HG00420. sv-channels-7samples: sv-channels trained on 7 samples (HG01053, HG01114, HG01881, HG02018, HG02924, HG03992, NA06991). sv-channels-LOCaS: sv-channels trained with the LOCaS strategy.

### sv-channels improves Mendelian concordance

One refers to Mendelian concordance when the variants in a trio are consistent with Mendelian inheritance. Variants in a child’s genome are concordant with Mendelian inheritance if they can be found in either the father’s or mother’s genome. To assess if sv-channels can improve Mendelian concordance of Manta DEL callsets, we run Manta on the 1KG sample HG00738 that belongs to a Puerto Rican child in a trio for which truth sets for the father (HG00736) and mother (HG00737) are available from the 1KG consortium. In the child’s truth set, the Mendelian concordance is 97,7%, which means that 97,7% of the DELs present in the child’s truth set are also present either in the father’s or in the mother’s truth set. The Mendelian concordance of the DEL callset of Manta is 53,2% (Figure 6). After filtering with sv-channels, the Mendelian concordance increases to 93.3%, which is close to the concordance reached in the child’s truth set.

**Figure 6.**
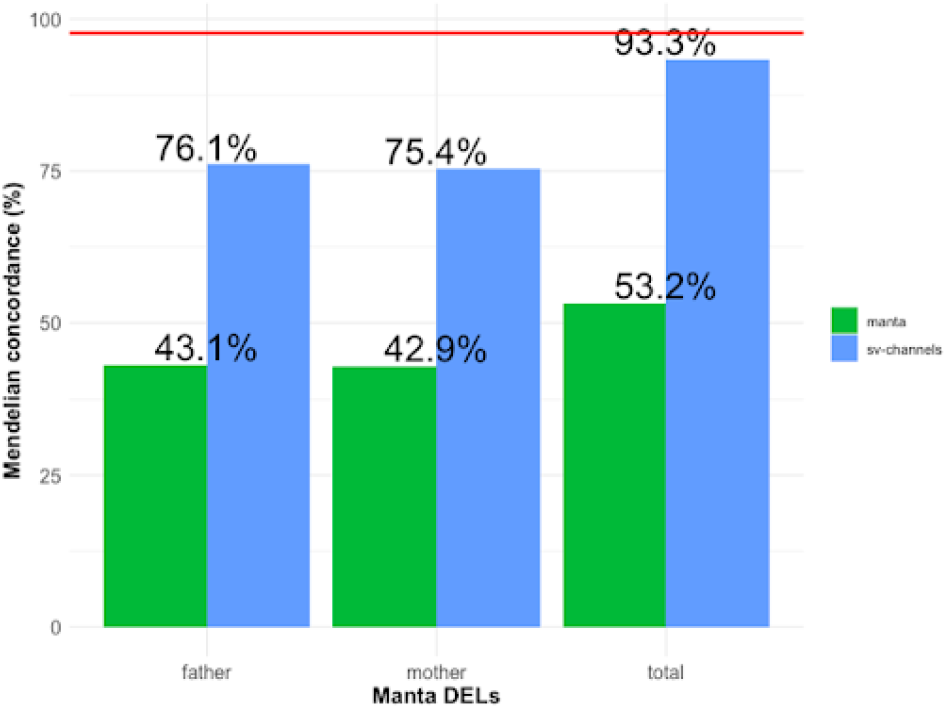
Mendelian concordance. (**A**) Mendelian concordance reported as percentage of Manta DELs that are called in the child (HG00738) that are also present in the father’s (HG00736) and mother’s (HG00737) truth sets, before (manta) and after the filtering with sv-channels (sv-channels). Bars show respectively when only the DELs shared between the child and the father are considered (father), only the DELs shared between the child and the mother (mother), and all the DELs shared between the child and either the mother or the father (total). The Mendelian concordance in the child’s truth set is shown as a red line.

## Discussion

We have introduced sv-channels, a novel deep learning approach aimed at removing false positive deletion calls from callsets generated by the SV-caller Manta.

We have shown its performance in several evaluations. First, when trained on a training set of seven samples from the 1KG dataset and applied on the hold-out test sample HG00420, the model showed an 82.87% increase in precision and a 10.54% decrease in recall with respect to the unfiltered Manta DELs callset. To assess the performance of the model in the absence of data leakage due to population deletions common between the training and the test set, we used two approaches (LOCaS and 2-fold LOCaS) where iteratively one chromosome at a time was used as test set in a nested cross-validation. The LOCaS approach showed a 74.08% increase in precision and a 15.91% decrease in recall with respect to the unfiltered Manta DEL callset, while the 2-fold LOCaS approach showed a 72.03% increase in precision and a 11.15% decrease in recall with respect to the unfiltered Manta DEL callset. We have also shown the performance of the model when using chromosome bins instead of whole chromosomes, using the LOBaS strategy (68.96% increase in precision and a 15.06% decrease in recall with respect to the unfiltered Manta DEL callset) and the 2-fold LOBaS approach (64.98% increase in precision and a 10.16% decrease in recall with respect to the unfiltered Manta DEL callset). This indicates that the channel representation is sufficient to improve the precision of Manta deletions.

Previous applications of Deep Learning for calling short variants (SNVs and indels, with DeepVariant) and long deletions (DeepSV) have seen the use of a high-dimensional tensor (image) representations for variant regions. Here we show that a one-dimensional representation is enough to distinguish true deletions from false positive calls.

The use of channel representation to filter other SV types (insertions, inversions, tandem duplications, and inter-chromosomal translocations) remains to be explored. Moreover, the performance of the model could be evaluated on the task of removing false positive calls in somatic SV callsets derived from the comparison of paired tumor and matched normal samples. In this case the input of the model would be formed by stacking the window-pair derived from the tumor sample on top of the window-pair derived from the matched normal sample. Considering that most genomic data can be naturally represented in one dimension in relation to the genomic position, the framework of sv-channels could be used to integrate additional channels with information, for instance, derived from evolutionary conservation, chromatin accessibility and long-range interactions in the genome. Including these channels in the model could help evaluate their importance in relation to the ability to separate false positive calls from true SVs. sv-channels could also be extended to read alignment data obtained from long read sequencing technologies such as those from PacBio and Oxford Nanopore Technologies (ONT). The sequences obtained with these technologies generate split reads when mapped to the genome at SV locations, so the channels containing split read information should be well suited to improve the precision of SV callers that use long read sequencing data.

## Acknowledgements

The project was supported by the Netherlands eScience Center, Grant ID: 027.016.G03, and by SURF, computing infrastructure grant number e-infra170206.

