## Supplemental Materials for "sv-channels: filtering genomic deletions using one-dimensional convolutional neural networks"

[1]Center for Molecular Medicine, University Medical Center Utrecht, Universiteitsweg 100, 3584 CG Utrecht, The Netherlands; [2]Oncode Institute, Office Jaarbeurs Innovation Mile (JIM), Jaarbeursplein 6, 3521 AL Utrecht, The Netherlands; [3]Netherlands eScience Center, Science Park 402, 1098 XH Amsterdam, The Netherlands; [4]current affiliation: JSR Life Sciences, NGS-AI division, Epalinges, Switzerland; [5]current affiliation: Interdisciplinary Centre for Security, Reliability and Trust (SnT), University of Luxembourg, 29 Av. J. F. Kennedy, Luxembourg, Luxembourg

### Supplemental materials and methods

#### A. Model

The model was implemented in TensorFlow v2.4.1. Model weights were initialized using “lecun_uniform” initialization. The model is a “Sequential” model with:

1. one BatchNormalization layer;
2. one or more Convolution1D layers (padding=’same’, ‘lecun_uniform’ initialization and l2 regularizer) followed by a BatchNormalization layer, ReLU activation and a Dropout layer;
3. one Dense layer (‘lecun_uniform’ initialization and l2 regularizer) followed by a BatchNormalization layer, ReLU activation and a Dropout layer;
4. one BatchNormalization layer followed by a SoftMax activation.

The model was compiled with categorical cross entropy loss and the Adam optimizer, and trained with batch size equal to 32 for 50 epochs.

#### B. Hyperparameter optimization

The following hyperparameters of the model were optimized:

1. number of convolutional filters (default value: 4);
2. number of convolutional layers (default value: 1);
3. size of the convolutional filters (default value: 7);
4. number of nodes in the Dense layer (default value: 4);
5. Dropout rate (default value: 0.2);
6. initial learning rate (default: 1e-4);
7. regularization rate (default: 1e-1).

Optimization was carried out using sequential Bayesian optimization with Gaussian processes using the gp_minimize function of the scikit-optimize library v0.9.0 (Head et al., 2018) with the negative expected improvement (‘EI’) as the acquisition function, using 50 iterations and selecting for negative mean balanced_accuracy as specified in the scikit-learn documentation across the test folds of the inner cross-validation step of the nested cross-validation.

#### C. Overlap among callsets

The overlap among the DEL callset (Manta, GRIDSS, sv-channels and truth sets) was assessed using the function countBreakpointOverlaps of the R Bioconductor package StructuralVariantAnnotation version 1.12.0 (Cameron et al., 2022) with the following parameter values:

- maxgap = 100
- sizemargin = 0.25
- ignore.strand = FALSE
- restrictMarginToSizeMultiple = 0.5
- countOnlyBest = TRUE

The SV type was inferred using the simpleEventType function provided in the GitHub repository of GRIDSS: https://github.com/PapenfussLab/gridss/blob/7b1fedfed32af9e03ed5c6863d368a821a4c699f/example/simple-event-annotation.R#L9

#### D. Simulated data

Read alignment data with simulated heterozygous and homozygous SVs were generated using the sv-gen workflow version v1.0.0 (Kuzniar and Santuari, 2023). We varied three parameters: read depth of coverage (coverage), insert size of the sequenced fragment (insert-size) and read length (read-length). For each parameter, we simulated 2,000 SVs of each type: deletions (DEL), insertions (INS), inversions (INV), tandem duplications (DUP) and inter-chromosomal translocations (TRA), for a total of 10,000 SVs inserted in chromosome 10 and chromosome 12 of the human reference genome hs37d5 of the 1000 Genomes project (ftp://ftp.1000genomes.ebi.ac.uk/vol1/ftp/technical/reference/phase2_reference_assembly_sequence/hs37d5.fa.gz). Simulated read alignment data was generated from the chromosomes containing the simulated SVs. The following parameters were varied: coverage (5x, 10x, 15x, 30x, 45x, 60x, 75x, 90x; with fixed insert-size at 500 bp and read-length at 150 bp), insert-size (200 bp, 250 bp , 300 bp, 400 bp, 500 bp, 600 bp; with fixed coverage at 30x and read-length at 150 bp) and read-length (36 bp, 50 bp, 75 bp, 100 bp, 150 bp, 250 bp; with fixed coverage at 30x and insert-size at 500 bp). The simulated SVs were called with Manta v1.1.0. Manta deletion (DEL) calls were filtered with sv-channels version v0.2.0 (Santuari et al., 2023) training the model using a 10-fold cross-validation. We used the exclusion list from ENCODE for the hg19 genome version downloaded from https://www.encodeproject.org/files/ENCFF001TDO/@@download/ENCFF001TDO.bed.gz to filter out SVs in problematic regions and we also filtered out SVs in regions containing the IUPAC ambiguity code N.

#### E. Training on the 1KG dataset

We selected a set of seven samples from the high-coverage 1KG dataset (Byrska-Bishop et al., 2022) for training: HG01053, HG01114, HG01881, HG02018, HG02924, HG03992, NA06991. 30% of the training set was used for validation. The trained model was tested on the sample HG00420 (Figure 5). The hyperparameters of the model were optimized using an inner 10-fold cross-validation, training the model for 50 epochs and using 50 iterations of the gp_optimize function. This resulted in the model called “sv-channels_model1”. In Figure 5 a nested Leave-One-Chromosome-and-Sample-Out Cross-Validation (LOCaS) strategy was used as described in the schema shown in Figure 3. The same seven samples specified before were used for training. The sample HG00420 was used as test sample in the outer cross-validation and sample HG01053 was used as test sample in the inner cross-validation. DELs in chromosome 22 of the sample HG00420 were used as validation set in the outer cross-validation and DELs in chromosome 22 of the sample HG01053 were used as validation set in the inner cross-validation. The same choice of training samples, test samples in the outer and inner cross-validations, and validation sets was used for the 2-fold LOCaS, the LOBaS and the 2-fold LOBaS strategies. For the LOBaS and 2-fold LOBaS strategies, the set of DELs for each chromosome was split into three sets of similar size (chromosome bins) based on the position of the first breakpoint and used in the nested cross-validation procedure instead of the whole chromosomes. To call SVs GRIDSS version v2.8.0 and Manta version v1.1.0 were used. To label Manta DELs as (true) DELs or noDELs the truth set file 1KGP_3202.Illumina_ensemble_callset.freeze_V1.vcf.gz (http://ftp.1000genomes.ebi.ac.uk/vol1/ftp/data_collections/1000G_2504_high_coverage/working/20210124.SV_Illumina_Integration/1KGP_3202.Illumina_ensemble_callset.freeze_V1.vcf.gz) was used, from which we extracted homozygous and heterozygous deletions for each sample using bcftools version 1.12 as follows:

VCF=1KGP_3202.Illumina_ensemble_callset.freeze_V1.vcf.gz; bcftools view -s $SAMPLE $VCF | bcftools view -i ’GT="alt"|GT="het"’ | bcftools view -i ’ALT="<DEL>"’ | bcftools view -i ’FILTER="PASS"’ > $SAMPLE.DEL.vcf

where the variable SAMPLE contains the sample name.

We used the exclusion list from ENCODE for the hg38 genome version downloaded from https://github.com/Boyle-Lab/Blacklist/blob/master/lists/hg38-blacklist.v2.bed.gz to filter out DELs in regions flagged as problematic by the ENCODE consortium (Amemiya et al., 2019) and we also filtered out DELs overlapping regions containing the IUPAC ambiguity code N.

#### F. Filtering deletions in a 1KG trio

We selected the samples from the following trio of the high-coverage 1KG dataset, a trio of Puerto Rican ethnicity:

1. Child: HG00738
2. Father: HG00736
3. Mother: HG00737

SVs were called with Manta v1.1.0. The truth set homozygous and heterozygous DEL calls were extracted from the 1KG truth set file 1KGP_3202.Illumina_ensemble_callset.freeze_V1.vcf.gz (http://ftp.1000genomes.ebi.ac.uk/vol1/ftp/data_collections/1000G_2504_high_coverage/working/20210124.SV_-Illumina_Integration/1KGP_3202.Illumina_ensemble_callset.freeze_V1.vcf.gz). The model “sv-channels_model1” was used to generate the sv-channels callset from the Manta callset. An SV quality (QUAL) threshold of 0.5 was used for the sv-channels callset.

#### G. Additional notes

The abstract’s readability was improved using the model GPT-4o.

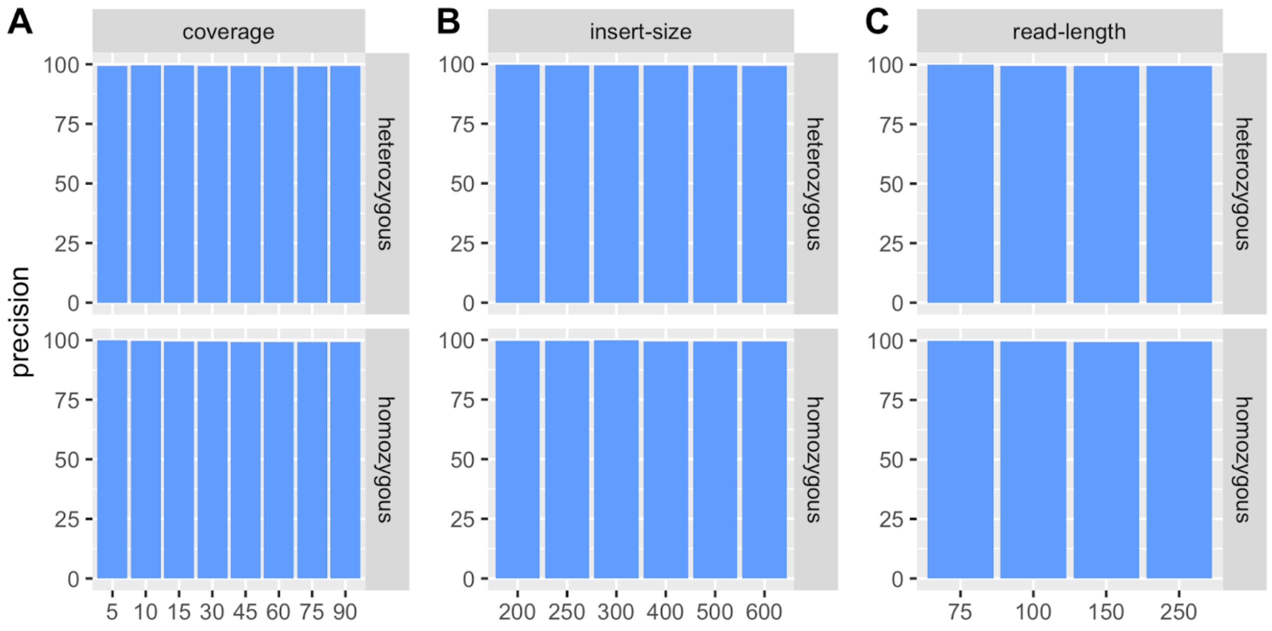


**Figure S1. Simulated data: precision**
Precision of sv-channels on simulated deletions at variable (**A**) coverage (**B**) insert size (**C**) read length.

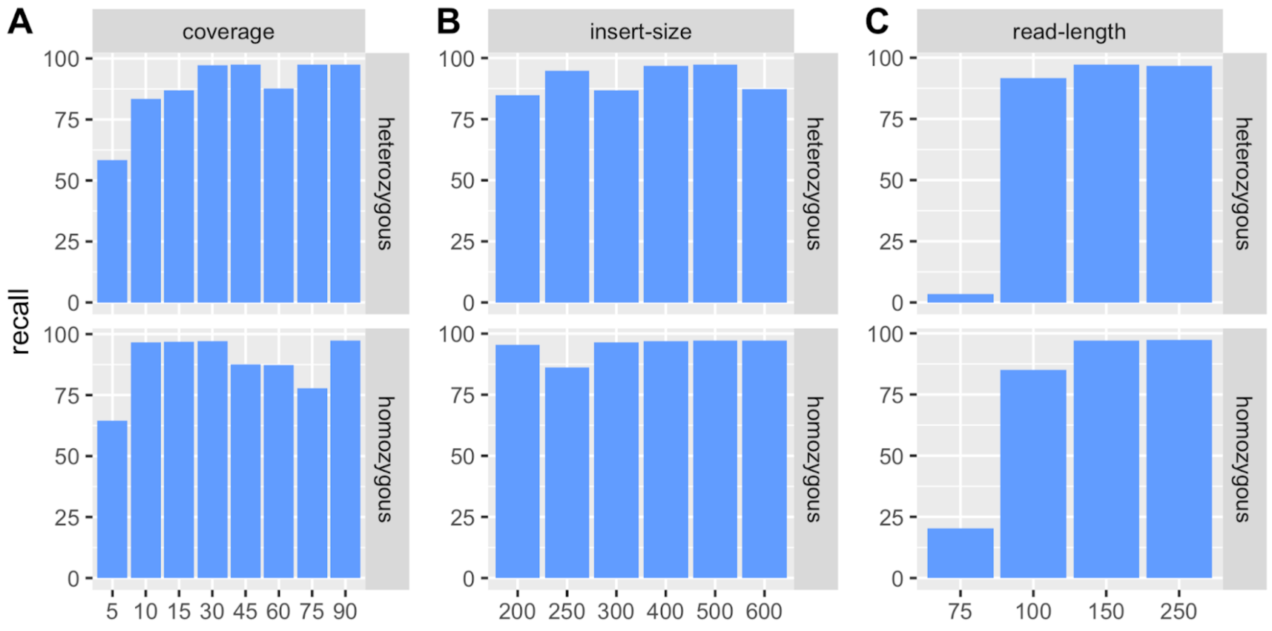


**Figure S2. Simulated data: recall**
Recall of sv-channels on simulated deletions at variable (**A**) coverage (**B**) insert size (**C**) read length.

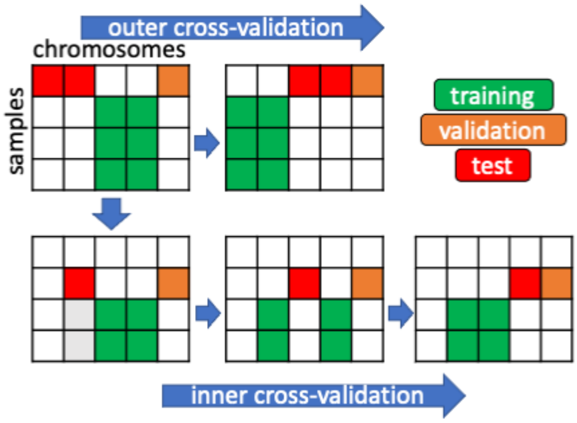


**Figure S3. Leave-One-Chromosome-and-Sample-out cross-validation**
Schema of the 2-fold nested Leave-One-Chromosome-and-Sample-out (2-fold LOCaS) cross-validation procedure. The inner cross-validation was used to optimize the hyperparameters using sequential Bayesian optimization with Gaussian Processes.

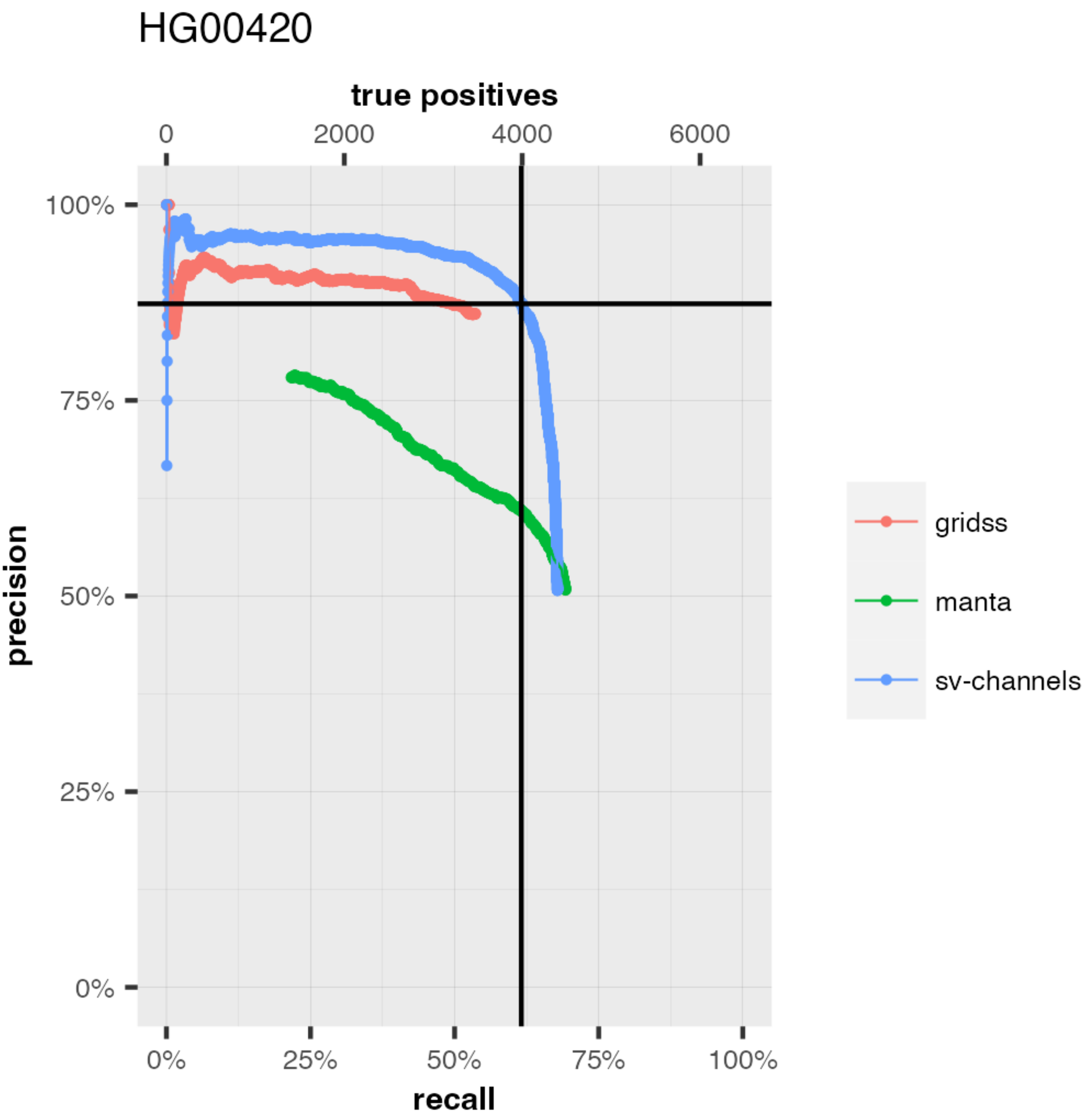


**Figure S4. 1KG dataset: 2-fold LOCaS**
Precision-Recall curves for sv-channels, GRIDSS and Manta. The sv-channels model was trained with 2-fold LOCaS using 7 samples for training (HG01053, HG01114, HG01881, HG02018, HG02924, HG03992, NA06991) and tested on sample HG00420. The intersection between the vertical line and the horizontal line indicates where the SV quality (QUAL) is equal to 0.5 for the sv-channels curve.

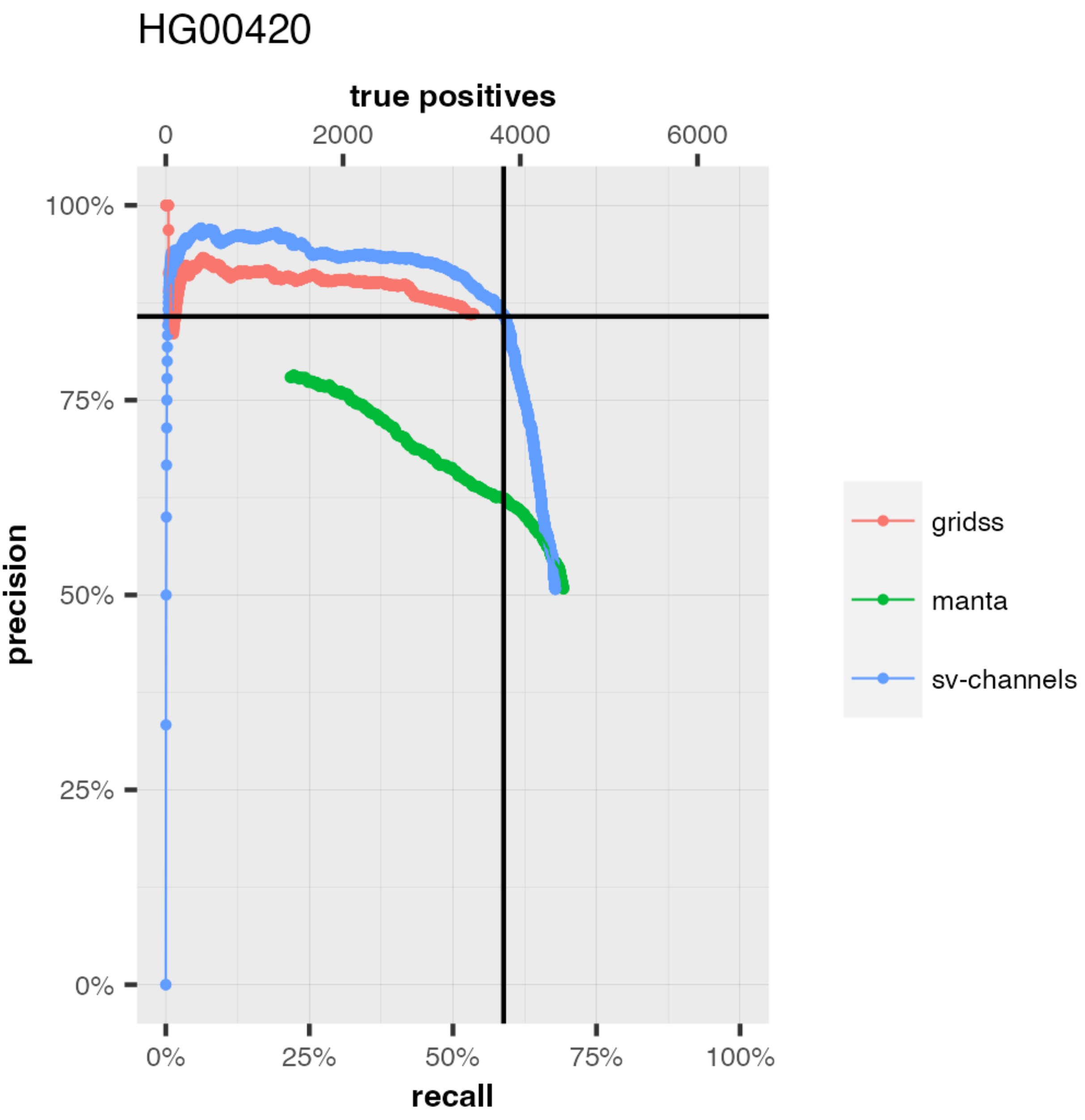


**Figure S5. 1KG dataset: LOBaS**
Precision-Recall curves for sv-channels, GRIDSS and Manta resulting from the LOBaS strategy (nested cross-validation with chromosome bins). The intersection between the vertical line and the horizontal line indicates where the SV quality (QUAL) is equal to 0.5 for the sv-channels curve.

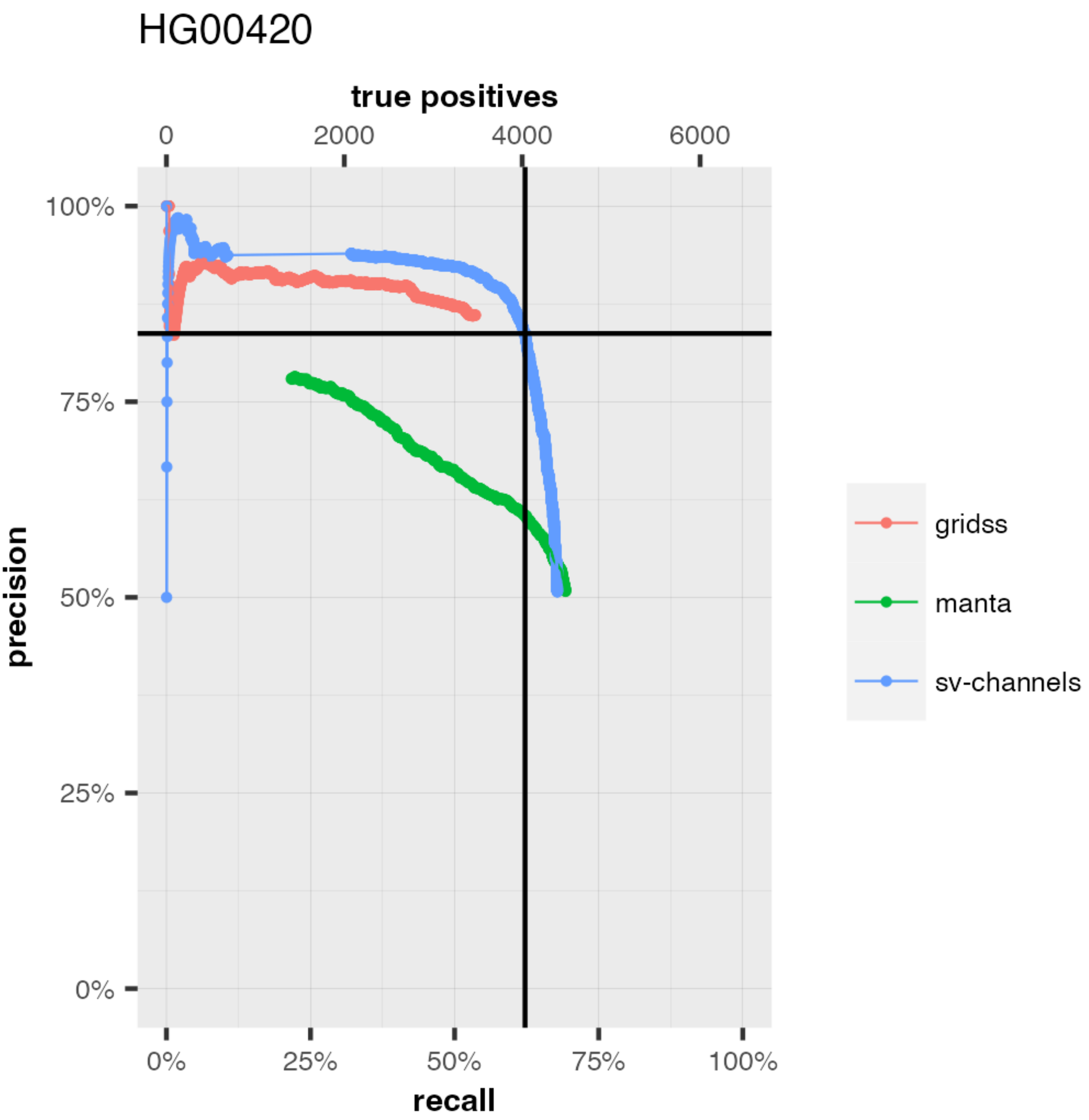


**Figure S6. 1KG dataset: 2-fold LOBaS**

Precision-recall curves for sv-channels, GRIDSS and Manta resulting from the 2-fold LOBaS strategy (2-fold nested cross-validation with chromosome bins). The intersection between the vertical line and the horizontal line indicates where the SV quality (QUAL) is equal to 0.5 for the sv-channels curve.

### Supplementary Tables

| Tool | Version | Wall clock time (hh:mm:ss) |
|---|---|---|
| GRIDSS2 without exclusion list | 2.8.0 | 05:04:36 |
| GRIDSS2 with exclusion list | 2.8.0 | 04:05:06 |
| Manta | 1.1.0 | 05:54:59 |
| svchannels extract-signals | 0.2.0 | 01:23:089 |
| svchannels generate-channels | 0.2.0 | 00:04:13 |
| svchannels score | 0.2.0 | 00:00:11 |

**Table S1. Runtimes**
Runtimes for GRIDSS2, Manta and sv-channels (commands “extract-signals”, “generate-channels” and “score”) when executed on a single core for the 1KG sample HG00420. The sv-channels command “extract-signals” is the most time-consuming command of sv-channels. Since it does not require the Manta output, it can be run in parallel with Manta.

| Number | Description |
|---|---|
| 1 | depth of coverage |
| 2 | one hot encoding for nucleotide A |
| 3 | one hot encoding for nucleotide C |
| 4 | one hot encoding for nucleotide T |
| 5 | one hot encoding for nucleotide G |
| 6 | one hot encoding for ’N’ |
| 7 | number of SOFT clipped reads on the LEFT in FWD direction |
| 8 | number of SOFT clipped reads on the RIGHT in FWD direction |
| 9 | number of SOFT clipped reads on the LEFT in REV direction |
| 10 | number of SOFT clipped reads on the RIGHT in REV direction |
| 11 | number of reads with a CIGAR "I" in FWD direction |
| 12 | number of reads with a CIGAR "I" in REV direction |
| 13 | number of reads with a number of mismatches (NM) equal to or greater than 11 and minimum mapping quality of 15 |
| 14 | number of reads SOFT clipped in both directions |
| 15 | number of reads with a CIGAR "D" in FWD direction |
| 16 | number of reads with a CIGAR "D" in REV direction |
| 17 | number of split reads with mapping quality lower than 10 |
| 18 | number of inter-chromosomal split reads |
| 19 | number of paired reads where the mate is unmapped |
| 20 | number of SPLIT reads with PLUS PLUS orientation |
| 21 | number of SPLIT reads with MINUS MINUS orientation |
| 22 | number of SPLIT reads with MINUS PLUS orientation |
| 23 | number of SPLIT reads with PLUS MINUS orientation |
| 24 | number of DISCORDANT reads with PLUS MINUS orientation |
| 25 | number of DISCORDANT reads with PLUS PLUS orientation |
| 26 | number of DISCORDANT reads with MINUS MINUS orientation |
| 27 | number of DISCORDANT reads with MINUS PLUS orientation |
| 28 | orphans: number of SPLIT reads with PLUS PLUS orientation |
| 29 | orphans: number of SPLIT reads with MINUS MINUS orientation |
| 30 | orphans: number of SPLIT reads with MINUS PLUS orientation |
| 31 | orphans: number of SPLIT reads with PLUS MINUS orientation |
| 32 | orphans: number of DISCORDANT reads with PLUS MINUS orientation |
| 33 | orphans: number of DISCORDANT reads with PLUS PLUS orientation |
| 34 | orphans: number of DISCORDANT reads with MINUS MINUS orientation |
| 35 | orphans: number of DISCORDANT reads with MINUS PLUS orientation |

**Table S2. List of channels**
List of channels used in sv-channels. The extension on each side of the breakpoints (half of window_size (Figure 1)) is set to 62 bp by default. buffer_size (Figure 1) is set to 8 bp by default. The channels with discordant reads (24,25,26,27,32,33,34,35) cover a region 8 times the window_size (62*2*8=992 bp for each window by default).

| Caller | total DELs | TP | FP | Precision (%) | Recall (%) | F1 score (%) |
|---|---|---|---|---|---|---|
| GRIDSS | 4086 | 3514 | 572 | 86 | 53.34 | 65.84 |
| Manta | 9008 | 4550 | 4458 | 50.51 | 69.06 | 58.35 |
| sv-channels | 4406 | 4070 | 336 | 92.37 | 61.78 | 74.04 |

**Table S3. Results for the 1KG sample HG00420**
This table shows the results when training the sv-channels model on seven 1KG samples (HG01053, HG01114, HG01881, HG02018, HG02924, HG03992, NA06991) and testing on sample HG00420. 30% of the training set was used as validation and the hyperparameters of the model were optimized using a 10-fold cross-validation. This table is relative to the experiment shown in Figure 5. The table shows name of the caller (caller), total number of deletions (Total DELs), number of true positives (TP), number of false positives (FP), precision, recall and F1 score for GRIDSS, Manta and sv-channels. For sv-channels, an SV quality (QUAL) threshold of 0.5 was used. For Manta and GRIDSS only good quality DEL calls ('PASS') were considered.

| Caller | total DELs | TP | FP | Precision (%) | Recall (%) | F1 score (%) |
|---|---|---|---|---|---|---|
| GRIDSS | 4036 | 3474 | 562 | 86.08 | 53.61 | 66.07 |
| Manta | 8840 | 4488 | 4352 | 50.77 | 69.26 | 58.59 |
| sv-channels LOCaS | 4270 | 3774 | 496 | 88.38 | 58.24 | 70.21 |
| sv-channels 2-fold LOCaS | 4566 | 3988 | 578 | 87.34 | 61.54 | 72.2 |
| sv-channels LOBaS | 4444 | 3812 | 632 | 85.78 | 58.83 | 69.79 |
| sv-channels 2-fold LOBaS | 4814 | 4032 | 782 | 83.76 | 62.22 | 71.4 |

**Table S4. Results for the 1KG sample HG00420: the LOCaS, 2-fold LOCaS, LOBaS and 2-fold LOBaS strategies.**
This table shows the results of sv-channels for the strategies LOCaS (Figure 3 and Figure 5), 2-fold LOCaS (Supplemental Figure 3 and 4), LOBaS (Supplemental Figure 5) and 2-fold LOBaS (Supplemental Figure 6) for the test sample HG00420, in comparison with Manta and GRIDSS. Column names are as described in Supplemental Table 3. Deletions for chromosome 22 of the test sample HG00420 were not considered because they were used as validation set.
